# Autophagy-associated Production of Antimicrobial Peptides hBD1 and LL37 Exhibits Anti-Bacillus Calmette-Guérin Effects in Lung Epithelial Cells

**DOI:** 10.1101/2020.02.21.959361

**Authors:** Rui-ning Wang, Hong-lin Liu, Yao-xin Chen, Qian Wen, Xin-ying Zhou, Jin-li Wang, Jia-hui Yang, Yan-fen Li, Zhen-yu Han, Li Ma

## Abstract

Antimicrobial peptides (AMPs) constitute important groups of bactericidal polypeptides against various microorganisms that exhibit their anti-bacteria activity through cleavage of precursor peptides into the active form of 50–100 amino acids in length. Various AMP cleavage mechanisms have been reported in different cell types; however, those in *Mycobacterium tuberculosis* (MTB)-infected lung epithelial cells remain unknown. In the present study, we found that MTB-infected lung epithelial cells expressed high level of the AMPs hBD1 and LL37 to kill intracellular MTB as the first-line immune barrier against MTB infection. Notably, their production in the lung epithelial cells was closely related to the function of autophagosomes and lysosomes. Experimental induction of autophagy in lung epithelial cells could enhance the expression of active hBD1 and LL37 at the post-transcriptional level, whereas silencing of these two active AMPs could decrease the bactericidal effect of autophagy. These findings indicated that cleavage of peptide precursors to form active AMPs might constitute a previously unrecognized antibacterial mechanism of autophagy.

**Author summary:** LM and RW conceived and designed the experiments; RW performed the experiments and analyzed the data; QW analyzed the data and contributed reagents/materials/analysis tools; HL performed the experiments; XZ, JY, YL and ZH analyzed the data. LM and RW drafted the manuscript.

## Introduction

Tuberculosis (TB), the second leading cause of fatal infectious disease, results from *Mycobacterium tuberculosis* (MTB) infection that mainly affects the lung. Until 2019, over 10.0 million people have been infected by MTB among whom 186,772 carried rifampicin-resistant MTB/multidrug-resistant [1]. As a barrier against infection, the alveolar epithelium constitutes an essential component of innate and mucosal immunity in the lung. Composed of type I and II pneumocytes, it can detect pathogens via pattern recognition receptors and secrete surfactant proteins, cytokines, chemotactic factors, and antimicrobial peptides (AMPs). MTB can invade alveolar epithelial cells and proliferate therein [2–4], eventually leading to host cell necrosis, tissue damage, and epithelial barrier damage [5]. However, although AMPs are known to play distinct roles in the barrier function against MTB invasion, they are relatively poorly characterized compared with other effector molecules.

AMPs comprise a group of polypeptides consisting of 50–100 amino acids that exist in many tissues and cell types, exhibit antimicrobial activity against a variety of microbes, such as bacteria, fungi, and some viruses including MTB [6], and can be divided into three families: defensins, cathelicidins, and histatins. Defensins can be further divided into α-, β-, and θ-defensins. α-defensins, primarily detected in neutrophils, are also termed human neutrophil peptides. β-defensins, mainly expressed in epithelial cells [7,8], exhibit wide taxonomic distribution [9,10]. Notably, MTB infection can stimulate human corneal fibroblasts [11], human endothelial cells [12], and human airway epithelia [13,14] to secrete β-defensin 1 (hBD1). Cathelicidins exhibit distinct structure and evolution from defensins albeit similar wide distribution and abundance [15]. LL37, the only type of human cathelicidin identified to date, is mainly expressed in neutrophils and respiratory epithelial cells. LL37 has important roles in various types of infectious disease and lung immune responses [16] including anti-TB function and can inhibit intracellular MTB growth [17].

Owing to the indispensable role of AMPs in the first-line defense barrier to MTB invasion, their expression mechanism has received considerable attention. AMPs are generally expressed as precursors and cleaved by various proteases to release the active peptides [18], with different mechanisms depending on the cell and AMP type. The human cathelicidin gene (*CAMP*) encodes an 18 kDa inactive precursor protein (hCAP18). In human neutrophils, the hCAP18 C-terminus is released by proteolytic hydrolysis, forming an active antimicrobial peptide consisting of 37 amino acids, termed LL37 [19]. hCAP18 can also be cleaved by elastase from azurophilic granules during the exocytosis and phagocytosis process in neutrophils [20]. Defensins also exhibit similar shear mechanism. α-defensins are generally synthesized in neutrophils and stored as the active peptides in granules after being processed from their precursors to the mature form [21–23]. Nevertheless, AMP processing mechanisms are also vary depending on microbial structures and inflammatory mediators [24,25] and remain incompletely understood.

Notably, it has been reported that autophagy contributes to AMP cleavage and anti-MTB activity. Generally, autophagy is identified as a physiological process of cells to maintain self-homeostasis by degrading intracellular contents through acidification and hydrolase via lysosome fusion [26]. Based on this degradation process, we hypothesized that autophagy also plays a role in cutting proteins. Consistent with this, autophagosomes can selectively encapsulate innocuous cytoplasmic components via autophagic adapter protein p62 and process them into short peptides with anti-MTB antimicrobial activity [27]. Moreover, autophagy can also exert anti-MTB activity through multiple mechanisms; in turn, MTB infection can induce autophagy in various cells. For example, macroautophagy participates in MTB clearance in mouse and human macrophage cell lines [26,28]. It is generally believed that autophagy can encapsulate MTB-containing phagosomes for lysosome fusion, wherein the wrapped MTB is degraded by proteolytic enzymes [29]. Furthermore, the ribosomal protein rps30 can be degraded by autophagy via p62 to produce short peptides, which act in concert with lysosomal hydrolases to kill MTB in autophagosomes [27].

Therefore, in this study we examined whether other novel AMPs might be produced by autophagic degradation and whether autophagy could cleave AMP precursors to active peptides as a new mechanism of autophagic anti-MTB functionality. We used A549 and BEAS-2B cells as epithelial cell models to screen for highly expressed AMPs following MTB infection, identifying AMPs hBD1 and LL37 and examining their roles in defense against MTB invasion in lung epithelial cells. To further explore the mechanism, we correlated the active peptide levels of hBD1 and LL37 with autophagy and autolysosome formation and verified the importance of AMPs in autophagy-mediated MTB lethality. Expounding these mechanisms will help to further understand the role of lung epithelial cells as the first-line barrier against MTB infection and the previously unrecognized mechanism of autophagy against intracellular MTB. Such knowledge may have considerable significance for augmenting anti-TB immune mechanisms and developing new TB treatment strategies.

## Materials and methods

### Patients and specimens

For the use of clinical materials for research purposes, prior approval was obtained from the Southern Medical University Institutional Board (Guangzhou, China). All tissue samples were collected and analyzed in a manner consistent with the prior written informed consent of the patients and healthy tissue donors. A total of three patients with lung TB were examined for lung pathology; TB was confirmed through positive acid-fast staining. Patient tissue was obtained during the puncture examination. Healthy tissue was obtained from a total of two volunteers as normal tissue adjacent to resected tumors. And the peripheral blood mononuclear cells (PBMCs) samples were collected and analyzed in a manner consistent with the prior written informed consent of the patients and healthy donors.

### Antibodies and reagents

The following reagents were used in this study: 3-methyladenine (3-MA; M9281; Sigma-Aldrich, St. Louis, MO, USA), dimethylsulfoxide (D2650; Sigma-Aldrich), bafilomycin A1 (Baf A1; sc-201550; Santa Cruz, Dallas, TX, USA), TRIzol reagent (15596-018; Invitrogen, Carlsbad, CA, USA), rapamycin (V900930; Sigma-Aldrich), and NH_4_Cl (A9434; Sigma-Aldrich). The following antibodies were used in this study: anti-GAPDH (TA-08; ZSGB-BIO, Peking, China), anti-ATG 5 (12994; Cell Signaling Technology, Danvers, MA, USA), anti-ATG 7 (8558; Cell Signaling Technology), anti-p62 (23214S; Cell Signaling Technology), hBD1 (ab170962; Abcam, London, UK), hCAP18 (ab80895; Abcam), and LL37 (ab69484; Abcam). The used antibody dilutions is 5% Bovine serum albumin dissolved in Tris Buffered Saline with Tween® 20 (TBST-10X) (A1933; Sigma-Aldrich).

### Bacterial and cell culture

*M. bovis* Bacillus Calmette-Guérin (BCG) strain 19015 was purchased from the American Type Culture Collection (ATCC, Manassas, VA, USA). BCG were grown in Middlebrook 7H9 broth medium (271310; Sparks, MD, USA) or on 7H10 agar medium (26271; Sparks) supplemented with BBL Middlebrook oleic acid-albumin-dextrose-catalase (OADC) in an incubator (37 °C, 5% CO_2_). Human monocyte-derived macrophages (HMDMs) was isolated from healthy donors’ blood and were cultured in Roswell Park Memorial Institute (RPMI) 1640 Medium supplemented with 10% fetal bovine serum (35-076-CV; Coring, New York, USA) and 20 ng/μL Recombinant Human GM-CSF (300-03; Peprotech, Rocky Hill, USA).BEAS-2B cells (CRL-9609; ATCC) were cultured in Bronchial Epithelial Cell Growth Medium (CM2010; CellCook, Guangzhou, China). THP1 cells (TIB-202; ATCC) were cultured in RPMI 1640 Medium supplemented with 10% fetal bovine serum (35-076-CV; Corning). A549 cells (CCL-185; ATCC) were cultured in Dulbecco’s modified Eagle’s medium supplemented with 10% fetal bovine serum. A549 cells were infected with GFP-LC3 lentivirus (17-10193; Millipore, Bedford, MA, USA) according to manufacturer instruction. After three weeks, stable transfectants expressing GFP were selected using fluorescence-activated cell sorting (BD Biosciences, San Jose, CA, USA). LC3 overexpression in the transfectants was confirmed by western blotting, and the formation of LC3 puncta in infected cells following stimulation with rapamycin was observed using fluorescence microscopy.

### siRNA and transient transfection

siRNAs were purchased from RiboBio (Guangzhou, China). A549 and BEAS-2B cells (at approximately 50% confluence) were transiently transfected with 40 pM si-control and siRNAs for other genes using Lipofectamine 2000 (Invitrogen) according to manufacturer instruction. siRNA sequences are listed in Table 1.

**Table 1.**
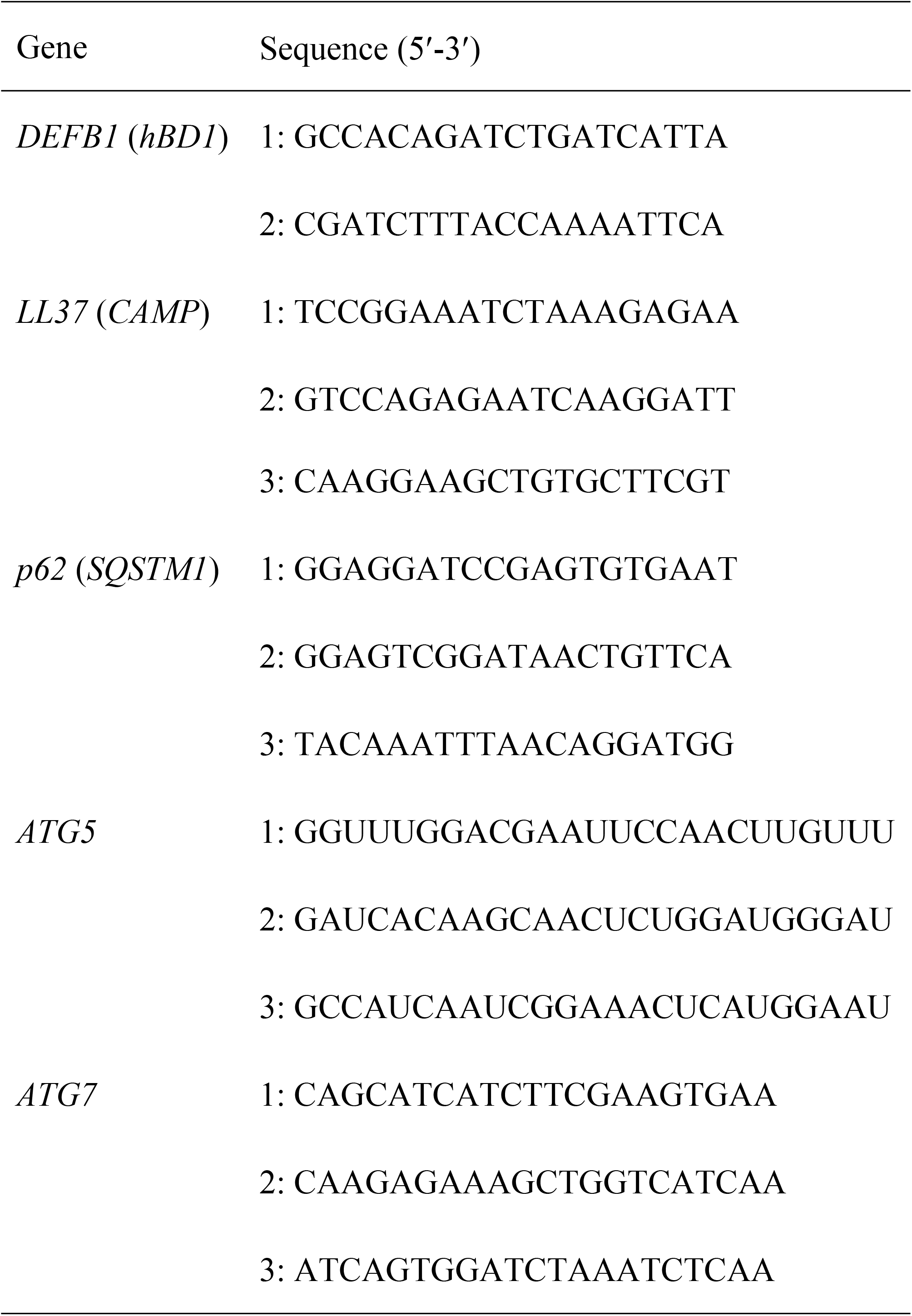
Sequence of siRNAs used in this study

### Western blotting

Whole cell extracts were prepared in the presence of protease inhibitor cocktail (Roche, Mannheim, Germany), phosSTOP phosphatase inhibitor cocktail (Roche), and 1 mM dithiothreitol (Biosharp, Shanghai, China). The cell extracts were resolved using sodium dodecyl sulfate-polyacrylamide gel electrophoresis, transferred to polyvinylidene fluoride membranes (Merck kGaA, Darmstadt, Germany), blocked using 5% bovine serum albumin for 1 h and incubated with diluted primary antibodies at 4 °C overnight. Western blots were developed using horseradish peroxidase-conjugated secondary antibodies, followed by detection with enhanced chemiluminescence. The results were analyzed using Adobe Photoshop CC 2017 (Adobe Systems Incorporated, San Jose, CA, USA); all the resultant bands were in the linear range (0–255 pixels).

### Confocal microscopy

A549 cells were grown on glass coverslips and treated with different stimulants, followed by fixation, permeabilization, and blocking. The coverslips were incubated with primary antibody at 4 °C overnight, then with secondary antibody for 1 h at room temperature. Nuclei were labeled with 4,6-diamidino-2-phenylindole staining. Before fixation, LysoTracker™ Blue DND-22 (Invitrogen, L7525) staining was performed by adding Lyso tracker (70 μM) to cells and incubating at 37 °C for 30 min. Coverslips were mounted with ProLong Gold antifade reagent (Invitrogen) and visualized using 60× mirror confocal microscopy (Zeiss, Göttingen, Germany).

### RNA isolation and quantitative real-time reverse transcription-polymerase chain reaction (qRT-PCR)

Total RNA was isolated using TRIzol reagent according to manufacturer recommendation. cDNA was synthesized using TransScript One-Step gDNA Removal and cDNA Synthesis SuperMix (AT311-03; TransGen Biotech, Guangzhou, China). For real-time qPCR analysis (40 cycles), TransStart Top Green qPCR SuperMix and an Eppendorf Mastercycler were used. *GAPDH* was used as housekeeping gene for data normalization. The primer sequences were: hBD1, 5′-TGAGATGGCCTCAGGTGGT-3′ (forward) and 5′-GCAGGCAGAATAGAGACATTGC-3′ (reverse); LL37, 5′-GTCCTCGGATGCTAA CCT CT-3′ (forward) and 5′-TCT GGT GAC TGC TGT GTC G-3′ (reverse); and *GAPDH*, 5′-GTC TCC TCT GAC TTC AAC AGC G-3′ (forward) and 5′-ACC ACC CTG TTG CTG TAG CCA A-3′ (reverse).

### Colony-forming units (CFU) assay

A549 and BEAS-2B cells were infected with BCG at a multiplicity-of-infection (MOI) of 5. After 1 h incubation at 37 °C, the infected cells were washed extensively with phosphate-buffered saline to remove extracellular mycobacteria, lysed using 1 mL of 0.01% Triton X-100 in distilled water, and 10-fold serially diluted to perform quantitative culture. The aliquots of each dilution were inoculated in triplicate on Middlebrook 7H10 agar plates supplemented with OADC. After 3-week incubation, the number of colonies was counted to determine the amount of intracellular viable bacilli. The survival rate of BCG at 72 h was calculated compared with that at 0 h.

### Co-immunoprecipitation (Co-IP)

Protein lysates used for Co-IP were prepared from A549 cells. Cells were harvested, lysed in IP buffer containing 0.3% CHAPS Detergent (10810118001; Roche), 40 mM HEPES (pH 7.5), 120 mM NaCl, 1 mM EDTA-2Na, and protease inhibitor cocktail (5892791001; Roche), then centrifuged at 12,000 x *g* (4 °C, 15 min). The supernatant was collected and incubated with p62 primary antibodies and protein G-agarose beads overnight at 4 °C. The beads were washed five times with IP buffer before processing for western blotting analysis.

### Immunohistochemistry

Immunohistochemistry staining was performed using a Dako Envision System (Carpinteria, CA, USA) following the manufacturer’s recommended protocol. Briefly, 4 μm thick tissue sections were incubated with anti-LL37 antibody at 4 °C overnight. Nuclear staining was regarded as a positive signal. Then, the tissue sections were incubated with biotinylated secondary antibody (Zymed, San Francisco, CA, USA), further treated with streptavidin-horseradish peroxidase complex (Zymed), immersed in 3-amino-9-ethyl carbazole, counterstained with 10% Mayer’s hematoxylin, dehydrated, mounted in Crystal Mount, and finally paraffin embedded. The tissue sections were scored by two independent observers.

### Electron microscopic analysis

A549 and BEAS-2B cells in T25 flasks were infected with BCG for 1 h with at a MOI of 5. At that time, the cells were fixed in 1% glutaraldehyde and processed for conventional transmission electron microscopy. Micrographs were taken from each sample, and the percentage of bacteria-containing cells were counted. For Electron microscopic analyzing, we thank the Central laboratory Southern Medical University, for technological assistance.

### Mass spectrometric analysis

To confirm the western blotting bands of AMPs, Mass spectrometric analysis was performed. A549 cells were infected with BCG for 48 h at a MOI of 5, samples were separated on 12% SDS-PAGE. The gel bands corresponding to the targeted protein were excised from the gel, reduced with 10 mM of DTT and alkylated with 55 mM iodoacetamide (I1149; Sigma-Aldrich). Then the gel was digested with trypsin (T1763; Sigma-Aldrich) at 37 °C for 6 h. Vortexing the gel for 5 min in 100% acetonitrile for 5 min and in 0.1% benzoic acid aqueous solution for 5 min, then in 100% acetonitrile for 5 min. The extractions were then centrifuged at 10,000 x *g* for 15 min. Then the sample were analyzed in Thermo Scientific™ Orbitrap Fusion™ Tribrid™, a liquid chromatography–mass spectrometry (LC–MS) system. And the data were analyzed by Proteome Discoverer software (1.2 version, Thermo Fisher Scientific, Waltham, MA, USA). For mass spectrometry analyzing, we thank the Central laboratory Southern Medical University, for technological assistance.

### Statistical analysis

All the measurement data are presented as the means ± standard deviation. Real-time PCR data were analyzed using one-way analysis of variance and least significant difference multiple comparison tests. Data between two groups were compared using the Student’s *t*-test. P-values were two-sided and a P-value < 0.05 indicated the presence of a statistically significant difference.

## Results

### Antimicrobial peptides hBD1 and LL37 are highly expressed in MTB-infected lung epithelial cells and exhibit anti-MTB activity

To determine whether AMPs play important roles in MTB infection in lung epithelial cells, we infected A549 and BEAS-2B cells with BCG and measured the mRNA levels of various AMPs. To confirm BCG infection, we performed electron microscopy (EM) analysis of BCG-infected A549 and BEAS-2B cells (S1 Fig). EM images showed that BCG (red arrow) were engulfed by A549 and BEAS-2B cells and enveloped by bilayer membrane autophagosomes (yellow arrow). However, as BCG enveloped by autophagosomes were destroyed, it was difficult to visualize the complete bacterial structure. This result illustrates that the mRNA levels of defensin hBD1 and cathelicidin LL37 were obviously higher than those of other AMPs in BCG-infected A549 cells (Fig 1A). Therefore, A549 cells were infected with BCG and LL37 and active hBD1 production was detected by qPCR or western blotting. BCG could effectively stimulate LL37 and hBD1 production at both mRNA and protein levels in A549 cells (Fig 1B, 1C&1D). To confirm these results, we detected LL37 and hBD1 expression in primary epithelial BEAS-2B cells, obtaining similar results (S2A, S2B&S2C Fig). Moreover, we found that hBD1 and LL37 mRNA levels were higher in peripheral blood mononuclear cells (PBMCs) from patients with MTB compared with healthy donors, indicating that these two AMPs might be involved in the process of MTB infection (Fig 1E&1F). To further confirm this result, we tested the level of LL37 by immunohistochemistry in lung biopsies of patients with MTB, revealing that their alveolar type II epithelial cells (but not immune cells) could produce much higher levels of LL37 than those in normal lung tissue sections and from patients with pneumoconiosis (Fig 1G). Similar results were obtained based on morphological features and hematoxylin staining (S3A&S3B Fig). This result was consistent with reports that AMPs are mainly expressed by epithelial cells [7,8]. Additionally, we detected *hBD1* and *LL37* mRNA levels in THP1 and HMDM cells compared to those in lung epithelial cells, revealing that both AMPs exhibited higher expression in lung epithelial cells, consistent with the immunohistochemistry results (S4 Fig) and further verifying that the choice of alveolar type II epithelial cells as the model in our study is reasonable because these cells are capable of producing high levels of AMPs [7,8].

**Figure 1.**
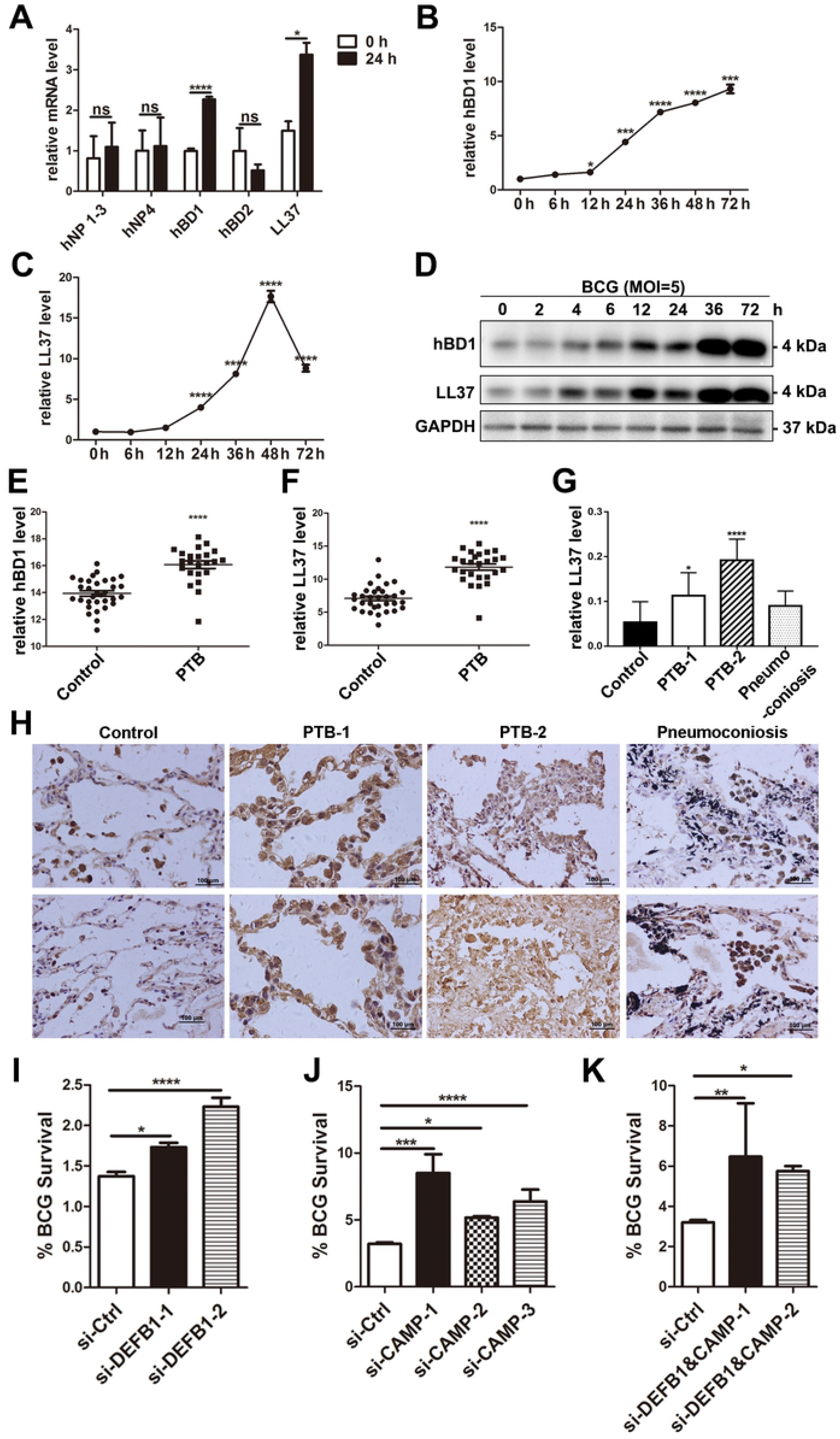
Antimicrobial peptides hBD1 and LL37 are highly expressed in MTB-infected lung epithelial cells and exhibit anti-MTB activity. (A) High level expression of hBD1 and LL37 was detected in A549 cells. The clls were infected with BCG for 48 h and the seven AMP mRNAs were evaluated by real-time PCR. (B&C) *hBD1* (B) and *LL37* (C) mRNA expression in BCG-infected A549 cells was detected at specified time points by real-time PCR. (D) The active peptides of hBD1 and LL37 were detected at specified time points and evaluated by western blotting. (E&F) High level expression of *hBD1* and *LL37* mRNA in PBMCs of patients with TB. (G) High level expression of LL37 was detected in human alveolar type II pneumocytes from patients with TB via immunohistochemistry. Micrograph shows strong LL37 immunostaining in human alveolar type II pneumocytes from patients with TB compared to that in healthy tissue donors and patients with pneumoconiosis (magnification x200). (H-J) Silencing of hBD1 or/and LL37 decreased intracellular BCG killing in A549 cells. The intracellular viable bacilli were determined by CFU assays at 72 h. Survival rate was calculated compared with that at 0 h. Data are expressed as the means ± standard deviation (s.d.). \**p* <0.05, \*\*\**p* < 0.001, \*\*\*\**p* < 0.0001. These experiments were performed independently at least thrice with similar results.

To confirm whether these two AMPs participated in the anti-MTB process in A549 and BEAS-2B cells, we performed CFU assays. A549 and BEAS-2B cells transiently transfected with si-control or si-DEFB1 or/and si-CAMP were infected with BCG. hBD1 or/and LL37 depletion significantly increased intracellular BCG survival in A549 (Fig 1I, 1J&1K) and BEAS-2B cells (S2G, S2H&S2I Fig). These results suggested that LL37 or/and hBD1 depletion impaired intracellular MTB elimination from lung epithelial cells, with these AMPs potentially functioning as effector molecules during the anti-MTB process in A549 and BEAS-2B cells.

### Autophagy influences hBD1 and LL37 active peptide levels

The autophagy-targeting molecule p62 (A170 or SQSTM1) is involved in killing MTB by specifically delivering ribosomal and bulk ubiquitinated cytosolic proteins to autolysosomes to produce AMPs [30]. Accordingly, we hypothesized that the autophagic level of cells might affect AMP production. To confirm our hypothesis, we used rapamycin, an autophagy agonist, to induce autophagy in A549 and BEAS-2B cells and measured hBD1 and LL37 production. BCG infection and rapamycin treatment increased autophagy level in A549 cells (Fig 2A). Rapamycin could effectively stimulate the production of active peptides of both AMPs, whereas the mRNA levels did not differ between rapamycin-treated and control groups (Fig 2A&2B). Similar results were obtained in BEAS-2B cells (S2 Fig). Furthermore, alternative autophagy stimulation in A549 cells via starvation culture medium with Earle’s balanced salt solution (EBSS) also increased AMP protein but not mRNA levels (Fig 2C&2D). Application of 3-MA to disturb the autophagic process in A549 and BEAS-2B cells decreased the active peptides of both AMPs whereas the mRNA levels remained unchanged (Fig 2A&2B and S2D, S2E&S2F Fig). Moreover, we used siRNA to silence autophagy-related protein 5 and 7 (ATG 5 and ATG 7) to generate autophagy-defective A549 cells. Notably, BCG could not stimulate active LL37 and hBD1 production in ATG 5- and ATG 7-deficient cells, despite increased mRNA levels (Fig 2E&2F). To further study the relationship between AMPs production and autophagy, we overexpressed hBD1 and LL37 precursor-mCherry and LC3-GFP in A549 cells. Upon BCG or rapamycin stimulation, hBD1 and LL37 precursor co-localization with LC3 puncta could be observed (Fig 3&4), supporting that AMP precursor was recruited into the autophagosomes. Conversely, under 3-MA treatment, hBD1/LL37 precursor and LC3 puncta co-localization was decreased (Fig 2G). These results showed that the autophagic level of lung epithelial cells affected hBD1 and LL37 active peptide but not mRNA levels and that AMP precursors entered autophagosomes, indicating that in the process of AMP production, autophagy plays an indispensable role at the post-transcriptional level.

**Figure 2.**
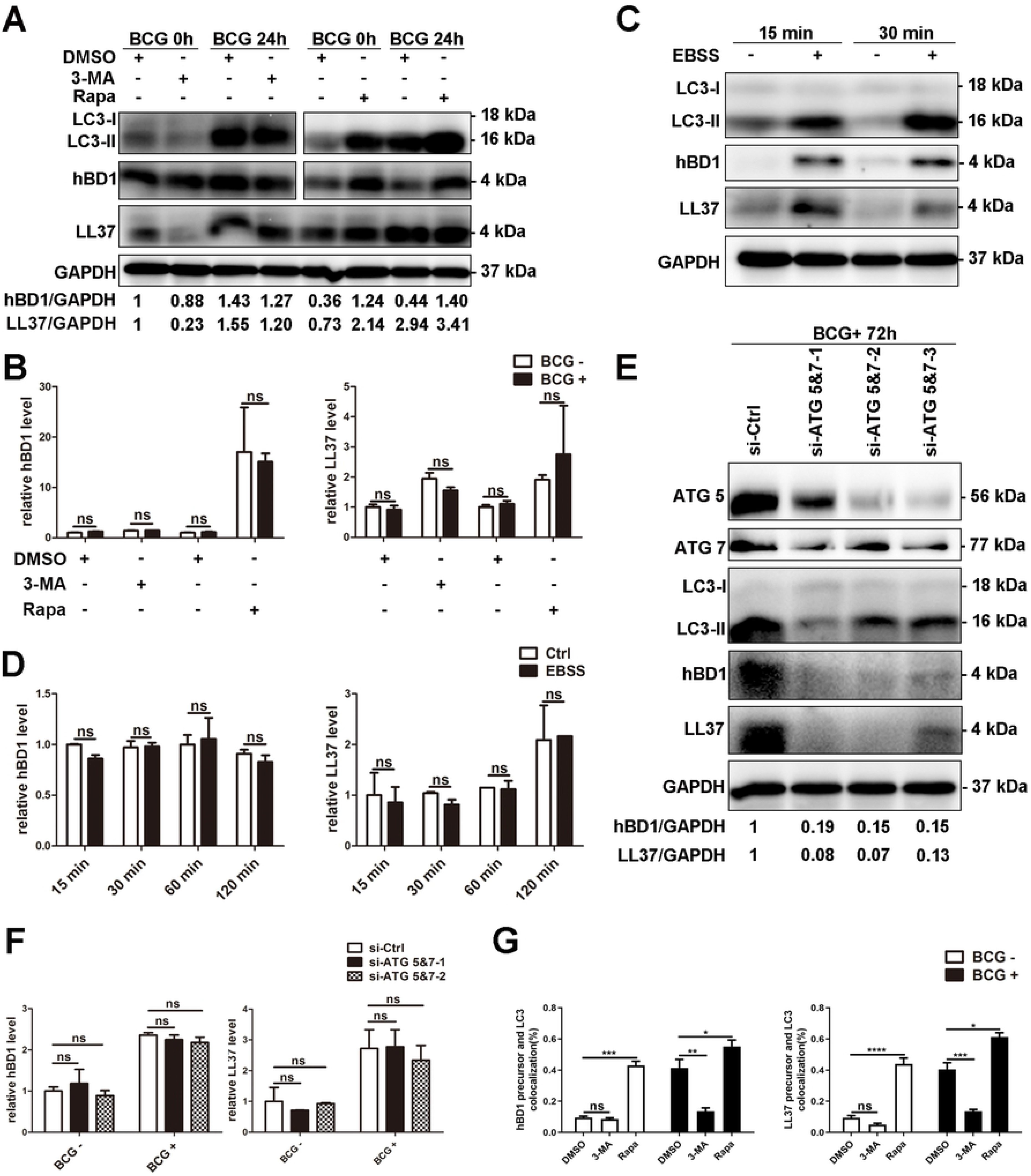
Autophagy influences the active peptide levels of hBD1 and LL37. (A) The autophagic level of A549 cells affected the active peptides level of hBD1 and LL37. A549 cells were pretreated with 4 μM rapamycin for 6 h and 5 mM 3-MA for 2 h and then infected with BCG for 24 h. The active peptide levels of hBD1 and LL37 were evaluated by western blotting. (B) The autophagic level of A549 cells did not affect the mRNA level of hBD1 and LL37. A549 cells were pretreated and infected with BCG as described above. The mRNA levels of hBD1 and LL37 were evaluated using real-time PCR. (C&D) Starvation-induced autophagy promoted the production of active hBD1 and LL37. A549 cells were cultured with EBSS culture medium at various time points. The expression of hBD1 and LL37 was detected by western blotting and real-time PCR. (E&F) Silencing ATG 5 and 7 disturbed the production of active hBD1 and LL37. A549 cells were transfected with 100 nM si-control or siRNA for ATG 5 and 7 for 72 h and infected with BCG. The expression of hBD1 and LL37 was detected by western blotting and real-time PCR. (G) The autophagic level of A549 cells affected the co-localization level of hBD1/LL37 precursor and autophagosomes. A549 cells stably expressing green fluorescent protein (GFP)– tagged LC3 (GFP-LC3) and mCherry Fluorescence Protein (mCherry)-tagged hBD1 or LL37 precursor (hBD1/LL37 precursor-mCherry) were pretreated with rapamycin and 3-MA and infected with BCG as described above. GFP-LC3 puncta (>1 μm) were observed and counted under confocal microscopy. Co-localization of hBD1/LL37 precursor and autophagosomes was detected by confocal microscopy. Data are expressed as the means ± standard deviation (s.d.) \**p* <0.05, \*\**p* <0.01, \*\*\**p* <0.001, \*\*\*\**p* <0.0001. These experiments were performed independently at least thrice with similar results.

**Figure 3.**
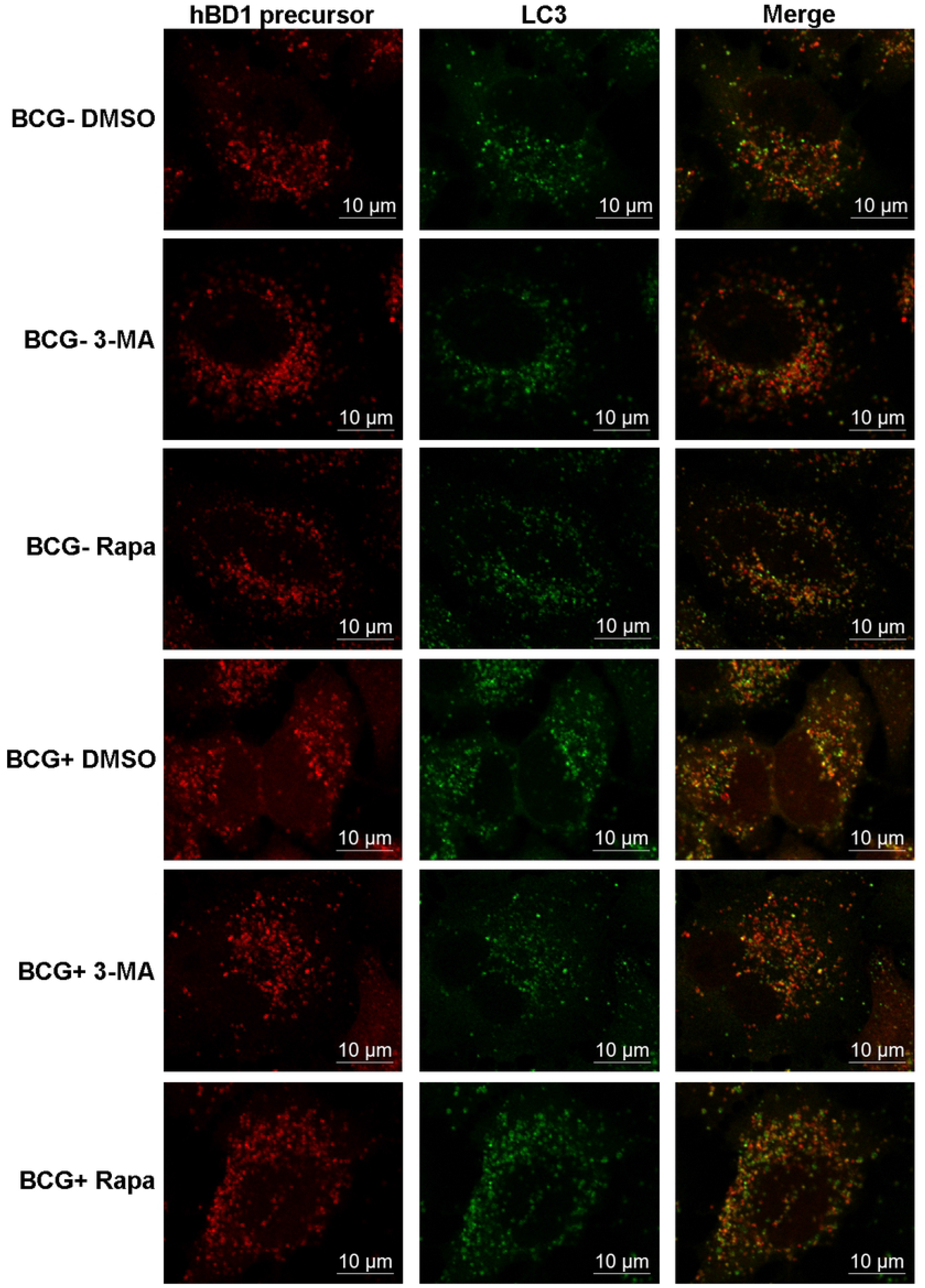
The autophagic level of A549 cells affected the co-localization level of hBD1 precursor and autophagosomes. A549 cells stably expressing green fluorescent protein (GFP)–tagged LC3 (GFP-LC3) and mCherry Fluorescence Protein (mCherry)-tagged hBD1 precursor (hBD1 precursor-mCherry) were pretreated with rapamycin and 3-MA as described above and infected with BCG as described above. GFP-LC3 puncta (>1 μm) were observed and counted under confocal microscopy. Co-localization of hBD1 precursor and autophagosomes was detected by confocal microscopy. These experiments were performed independently at least thrice with similar results.

**Figure 4.**
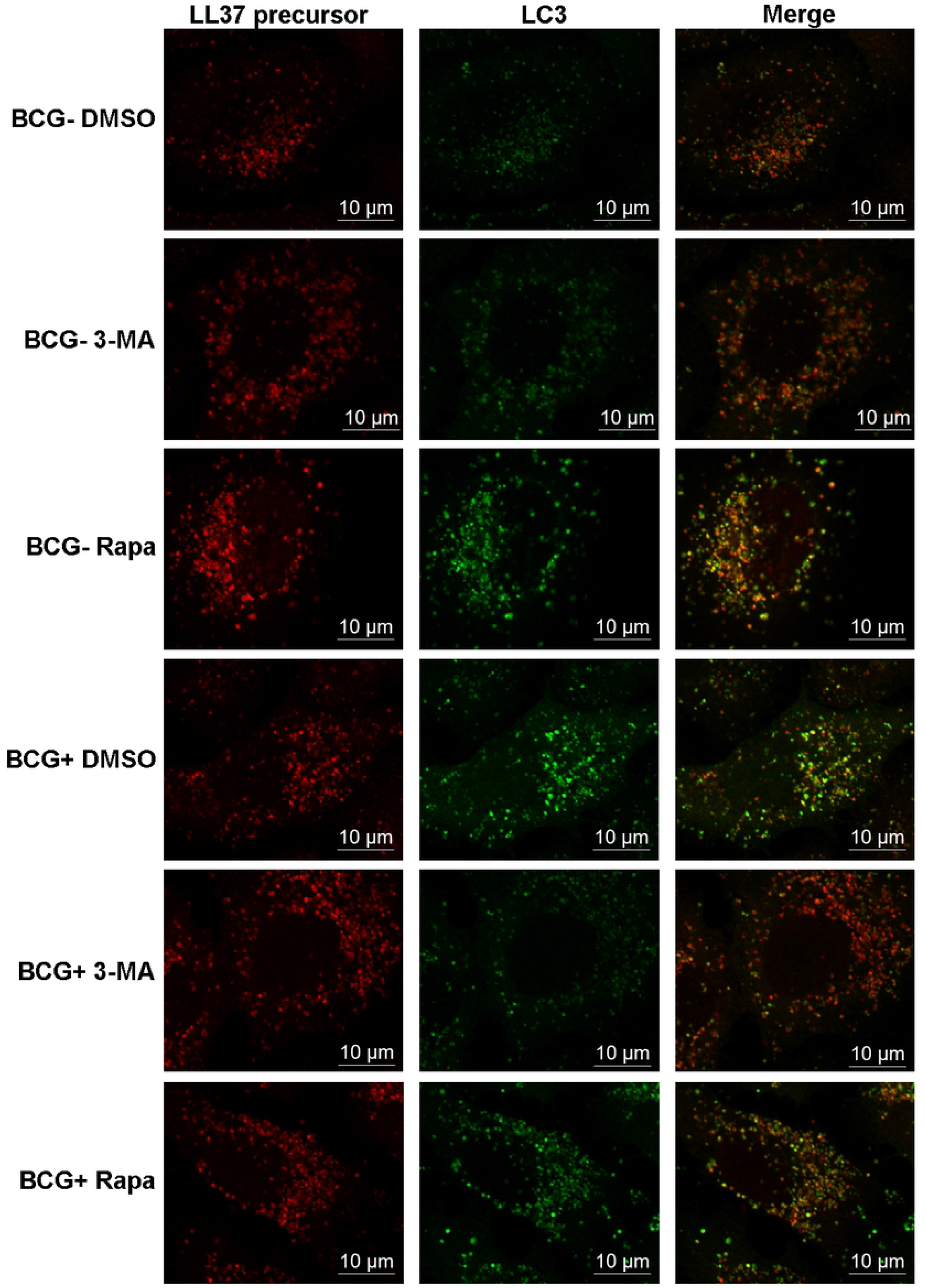
The autophagic level of A549 cells affected the co-localization level of LL37 precursor and autophagosomes. A549 cells stably expressing green fluorescent protein (GFP)–tagged LC3 (GFP-LC3) and mCherry Fluorescence Protein (mCherry)-tagged LL37 precursor (LL37 precursor-mCherry) were pretreated with rapamycin and 3-MA as described above and infected with BCG as described above. GFP-LC3 puncta (>1 μm) were observed and counted under confocal microscopy. Co-localization of LL37 precursor and autophagosomes was detected by confocal microscopy. These experiments were performed independently at least thrice with similar results.

### hBD1 and LL37 production are affected by lysosome function

To ensure complete sequestration during autophagy, the macromolecules and organelles should be delivered from autophagosomes to lysosomes, finally forming an autolysosome [31]. Thus, we hypothesized that the precursors of hBD1 and LL37 might be captured by autophagosomes and delivered to lysosomes to be cut into AMPs and exhibit their anti-MTB activity. We used Baf A1, an inhibitor of autophagosome formation that can block the fusion between autophagosomes and lysosomes, to treat BCG-infected A549 cells and measured the production of hBD1 and LL37 active peptides and mRNA. Baf A1 could effectively reduce active hBD1 and LL37 production (Fig 4A) whereas it did not affect their mRNA levels (Fig 4B). We next applied NH_4_Cl to disturb lysosomal and autophagosomal function, which obviously reduced active AMPs production (Fig 4C) but did not affect their mRNA levels (Fig 4D).

Furthermore, immunofluorescence to reflect the co-location of hBD1/LL37 precursor with autophagosomes and lysosomes showed that in A549 cells, the precursors of hCAP18 exhibited colocalization with autophagosomes and lysosomes, suggesting that the AMP precursors entered in autophagosomes (Fig 6&7). However, BCG infection of A549 cells treated with Baf A1 or NH_4_Cl did not alter the colocalization among hCAP18, autophagosomes, and lysosomes, even upon rapamycin stimulation (Fig 5E&5F). This indicated that autolysosome formation and function were not involved in AMP precursor recruitment. Thus, whereas the fusion of autophagosomes with lysosomes and lysosomal function are vital for active AMP production, these exert no effect on AMP precursor recruitment.

**Figure 5.**
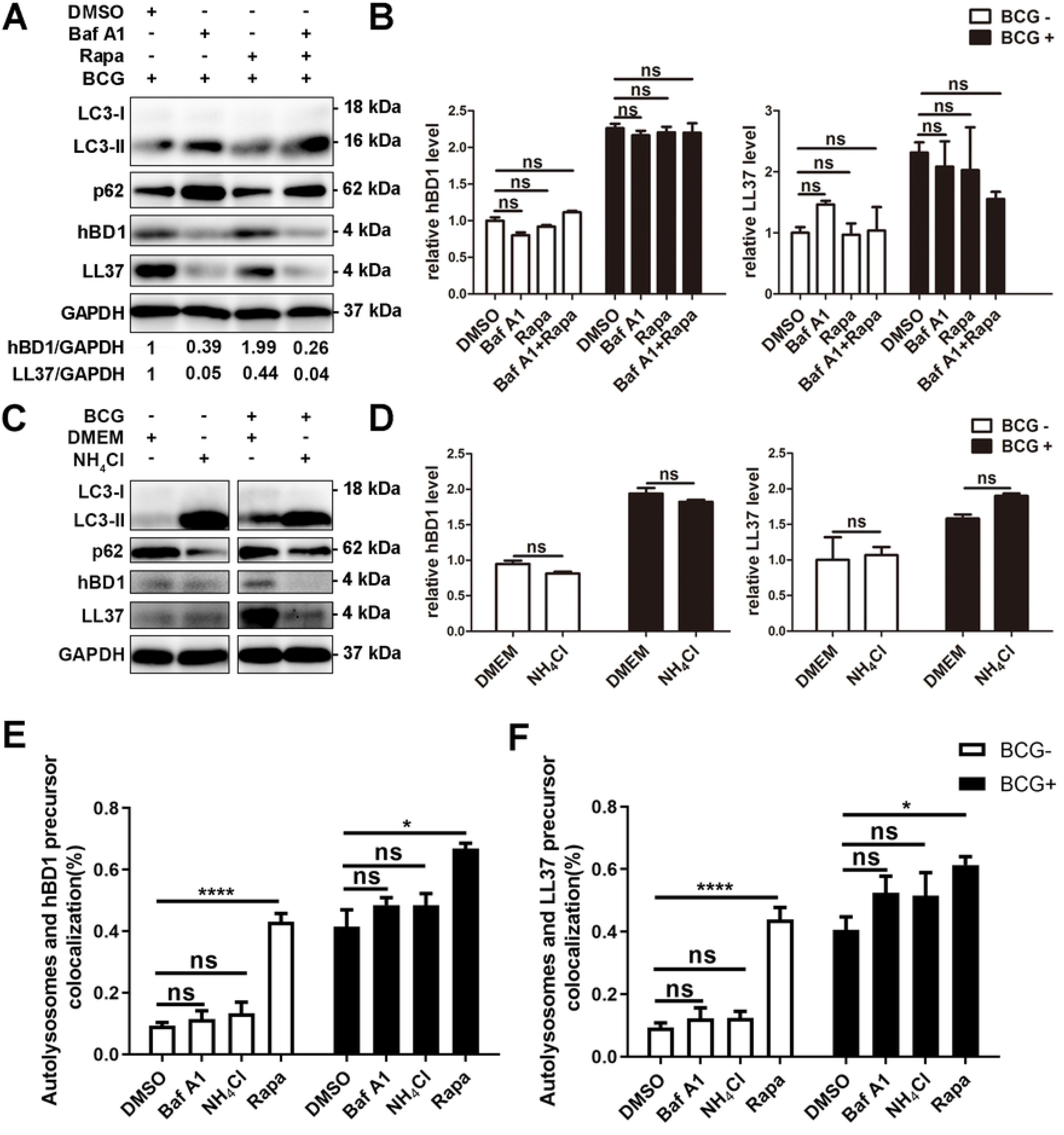
hBD1 and LL37 production was affected by the function of lysosomes. (A) The formation of autolysosomes affected the active peptide levels of hBD1 and LL37. A549 cells were pretreated with 0.1 mM Baf A1 or/and 4 μM rapamycin and infected with BCG for 48 h. The active peptide levels of hBD1 and LL37 were evaluated using western blotting. (B) The formation of autolysosomes did not affect the mRNA level of hBD1 and LL37. A549 cells were pretreated and infected with BCG as described above. The mRNA levels of hBD1 and LL37 were evaluated using real-time PCR. (C) The function of lysosomes affected the active peptide levels of hBD1 and LL37. A549 cells were pretreated with 10 mg/mL NH4Cl and infected with BCG for 48 h. The active peptide levels of hBD1 and LL37 were evaluated by western blotting. (D) The function of lysosomes did not affect the mRNA level of hBD1 and LL37. A549 cells were pretreated and infected with BCG as described above. The mRNA levels of hBD1 and LL37 were evaluated using real-time PCR. (E&F) The formation of autolysosomes did not affect the co-localization rates between hBD1/LL37 precursor and autolysosomes. A549 cells stably expressing GFP–tagged LC3 (GFP-LC3) and mCherry Fluorescence Protein (mCherry)-tagged hBD1 or LL37 precursor (hBD1/LL37 precursor-mCherry) were pretreated and infected with BCG as described as (A). Lysosome was labelled with lyso tracker, GFP-LC3 (>1 μm) and lyso tracker double positive puncta were determined as autolysosomes. Co-localization of hBD1/LL37 precursor and autolysosomes was detected by confocal microscopy. Data are expressed as the means ± standard deviation (s.d.) \**p* < 0.05, \*\*\*\**p* < 0.0001. These experiments were performed independently at least thrice with similar results.

**Figure 6.**
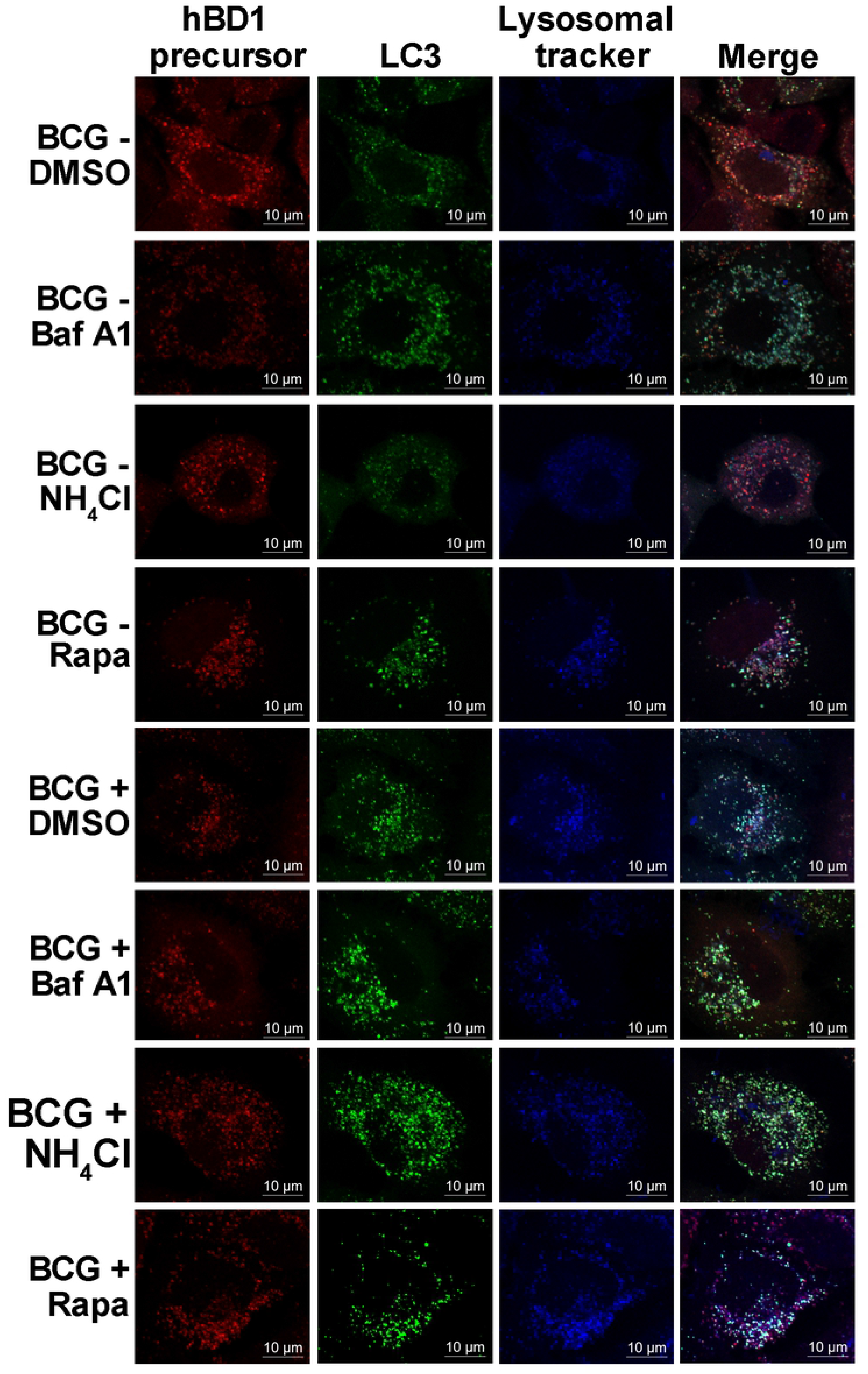
The formation of autolysosomes could not affect the co-localization rates between hBD1 precursor and autolysosomes. A549 cells stably expressing GFP–tagged LC3 (GFP-LC3) and mCherry Fluorescence Protein (mCherry)-tagged hBD1 precursor (hBD1 precursor-mCherry) were pretreated with 0.1 mM Baf A1 or/and 4 μM rapamycin and infected with BCG for 48 h. Lysosome was labelled with lyso tracker, GFP-LC3 (>1 μm) and lyso tracker double positive puncta were determined as autolysosomes. The function of lysosomes could not affect the co-localization rates between hBD1 precursor and autolysosomes. These experiments were performed independently at least thrice with similar results.

**Figure 7.**
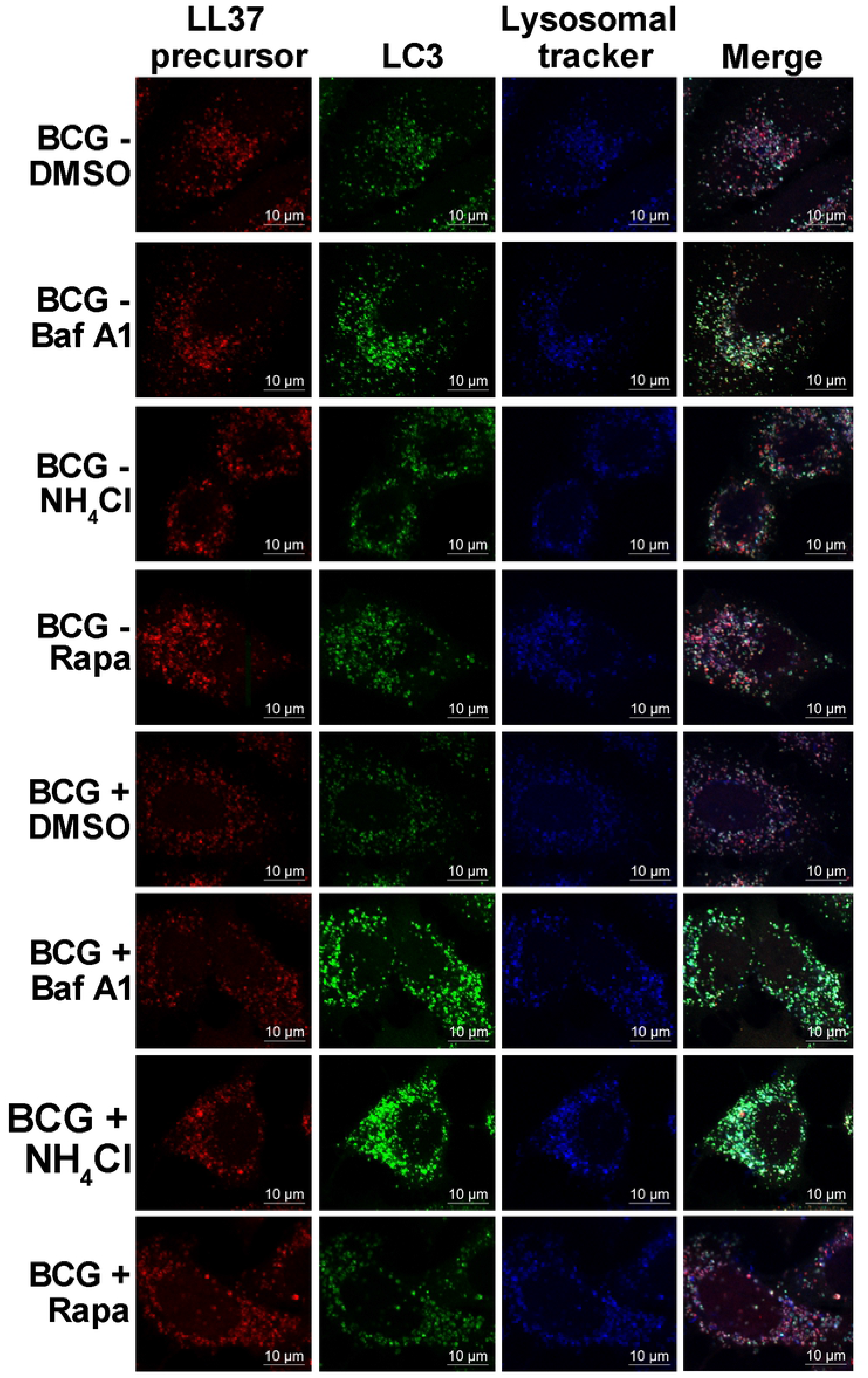
The formation of autolysosomes could not affect the co-localization rates between LL37 precursor and autolysosomes. A549 cells stably expressing GFP–tagged LC3 (GFP-LC3) and mCherry Fluorescence Protein (mCherry)-tagged LL37 precursor (LL37 precursor-mCherry) were pretreated with 0.1 mM Baf A1 or/and 4 μM rapamycin and infected with BCG for 48 h. Lysosome was labelled with lyso tracker, GFP-LC3 (>1 μm) and lyso tracker double positive puncta were determined as autolysosomes. The function of lysosomes could not affect the co-localization rates between LL37 precursor and autolysosomes. These experiments were performed independently at least thrice with similar results.

### Autophagy adapter protein p62 controls active AMP production by interacting with AMP precursors

p62/SQSTM1 is a kind of scaffold protein exhibiting multiple biological functions such as being a selective autophagy receptor for degradation by sensing ubiquitinated cargos [30]. We hypothesized that AMP precursors might be recruited into autophagosomes by selectively interacting with p62. To investigate whether p62 could influence AMP production, A549 cells transiently transfected with si-control or si-SQSTM1 were infected with BCG. Our analysis showed that endogenous p62 expression was effectively inhibited by the siRNA and that p62 depletion significantly reduced active hBD1 and LL37 but not mRNA production at indicated time points (Fig 8A&8B).

**Figure 8.**
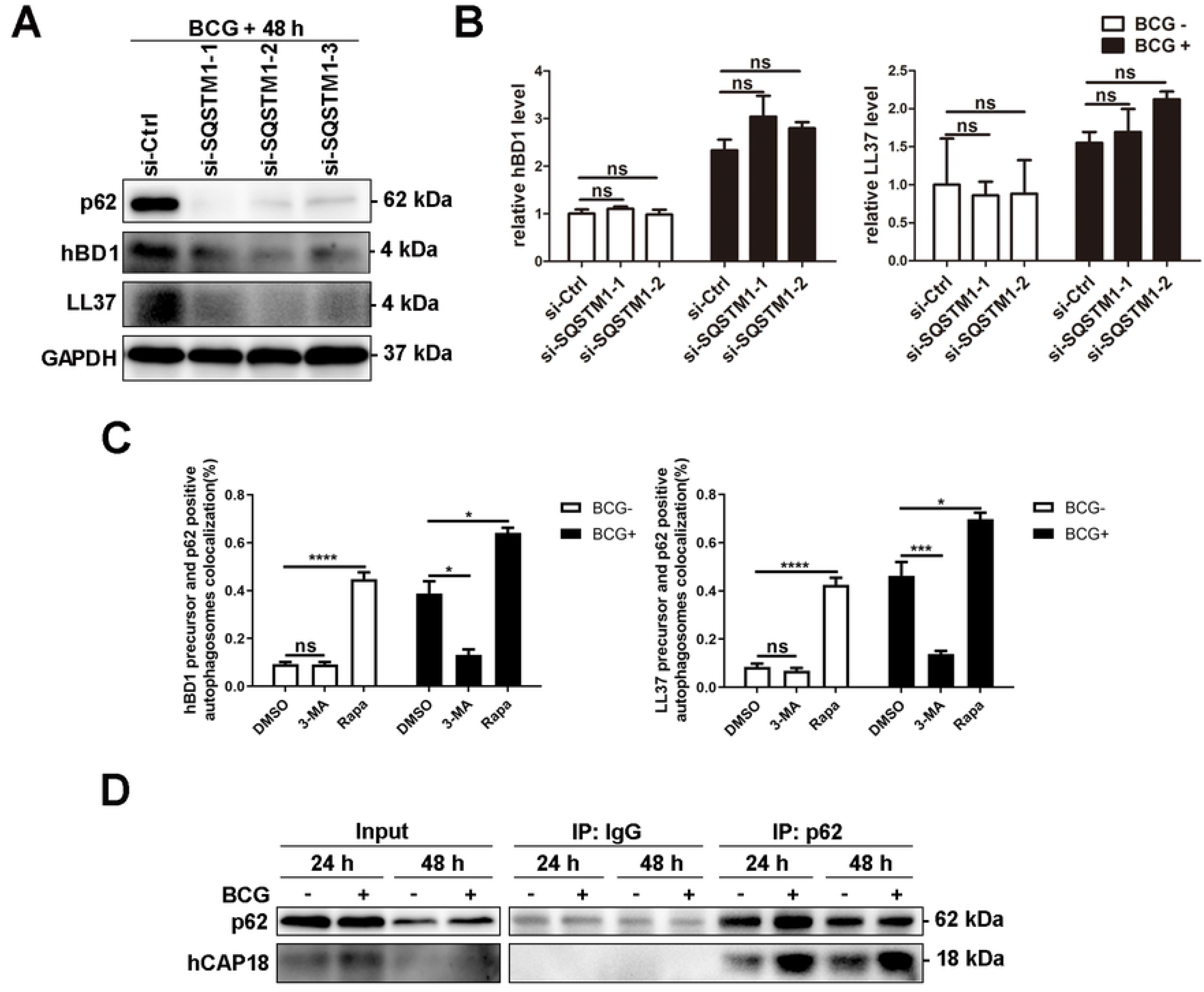
Autophagy adapter protein p62 controls the production of active AMPs by interacting with AMP precursors. (A&B) Silencing P62 affected the active peptide levels of hBD1 and LL37. A549 cells were transiently transfected with si-control or si-SQSTM1 to silence P62 and then infected with BCG. The expression of hBD1 and LL37 was detected using western blotting and real-time PCR. (C) The autophagic level of A549 cells affected the co-localization level of hBD1/LL37 precursor and p62-positive autophagosomes. A549 cells stably expressing green fluorescent protein (GFP)-tagged LC3 (GFP-LC3) and mCherry Fluorescence Protein (mCherry)-tagged LL37 precursor (LL37 precursor-mCherry) were pretreated with 3-MA and rapamycin and infected with BCG as described above. GFP-LC3 puncta (>1 μm) were observed and counted under confocal microscopy. Co-localization of hBD1/LL37 precursor, p62, and GFP-LC3, labeled using Alexa Fluor 647-coupled antibody against p62, was detected by confocal microscopy. (D) Direct interaction could be observed between p62 and hCAP18. A549 cells were infected with BCG for 24 and 48 h. The interaction between p62 and hCAP18 was detected by Co-IP with anti-p62 antibody followed by western blotting with anti-p62 and anti-hCAP18 antibodies. Data are expressed as the means ± standard deviation (s.d.) \**p* <0.05, \*\*\**p* <0.001, \*\*\*\**p* <0.0001. These experiments were performed independently at least thrice with similar results.

Additionally, p62, hBD1/LL37 precursor, and LC3 puncta co-localization was observed under BCG stimulation in A549 cells. The results revealed co-localization between p62 and hBD1/LL37 precursor in the cytoplasm and autophagosomes and indicated that the co-localization rates between hBD1/LL37 precursor and p62-positive autophagosomes were influenced by the autophagy level of A549 cells (Fig 8C, Fig 9&10). Furthermore, we performed Co-IP to directly measure and further confirm the interaction of p62 with AMP precursors. Under BCG stimulation, p62 interacted with hCAP18 (Fig 8D), suggesting that p62 is required for the autophagic production of active AMPs.

**Figure 9.**
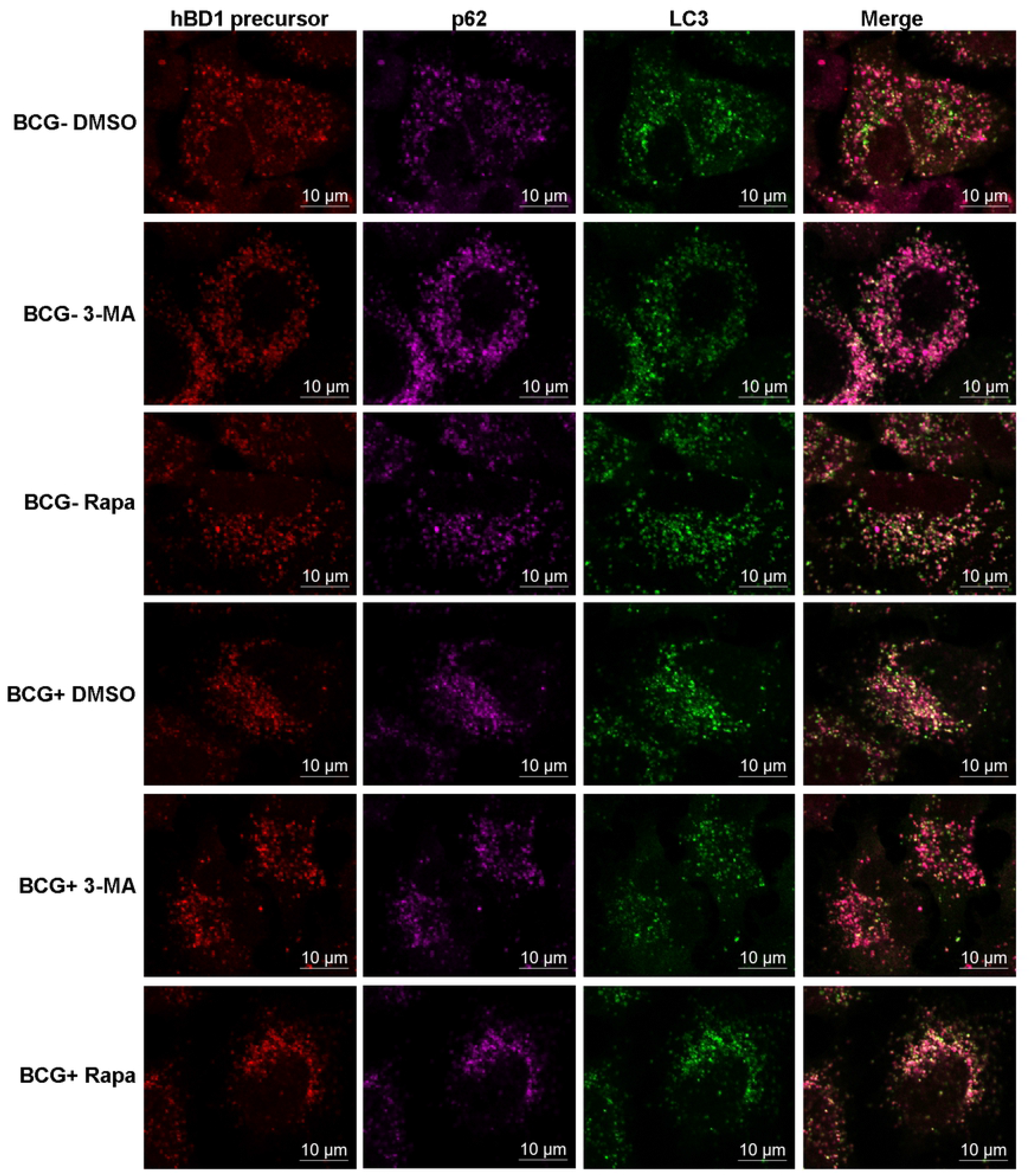
The autophagic level of A549 cells affected the co-localization level of hBD1 precursor and p62 positive autophagosomes. A549 cells stably expressing green fluorescent protein (GFP)-tagged LC3 (GFP-LC3) and mCherry Fluorescence Protein (mCherry)-tagged hBD1 precursor (hBD1 precursor-mCherry) were pretreated with 3-MA and Rapamycin and infected with BCG as described above. GFP-LC3 puncta (>1 μm) were observed and counted under confocal microscopy. Co-localization of hBD1, p62 and GFP-LC3, marked with Alexa Fluor 647-coupled antibody against p62, was detected by confocal microscopy. These experiments were performed independently at least thrice with similar results.

**Figure 10.**
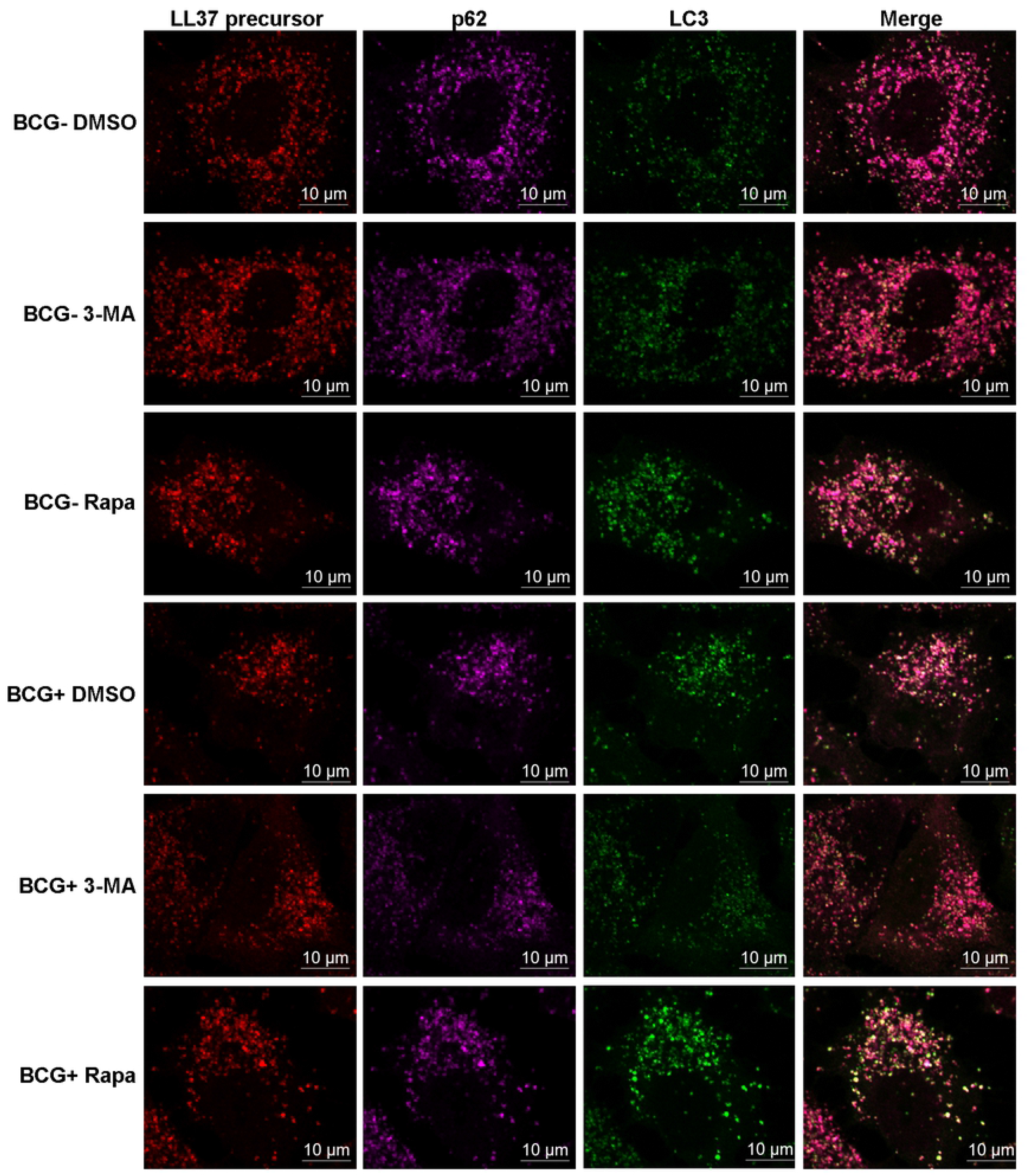
The autophagic level of A549 cells affected the co-localization level of LL37 precursor and p62 positive autophagosomes. A549 cells stably expressing green fluorescent protein (GFP)-tagged LC3 (GFP-LC3) and mCherry Fluorescence Protein (mCherry)-tagged LL37 precursor (LL37 precursor-mCherry) were pretreated with 3-MA and Rapamycin and infected with BCG as described above. GFP-LC3 puncta (>1 μm) were observed and counted under confocal microscopy. Co-localization of LL37, p62 and GFP-LC3, marked with Alexa Fluor 647-coupled antibody against p62, was detected by confocal microscopy. These experiments were performed independently at least thrice with similar results.

### Autophagy-related AMP production is a novel mechanism of autophagy-mediated MTB killing

To verify whether the post-transcriptional promotion of active AMP production by autophagy can play a role in killing intracellular MTB, a CFU assay was performed. Firstly, A549 and BEAS-2B cells were transiently transfected with si-control, si-DEFB1, or si-CAMP to silence hBD1 or/and LL37 and infected with BCG. Then, autophagy was further induced by rapamycin, and cells were lysed after 72 h. The results of CFU showed that under the induction of autophagy by rapamycin in A549 and BEAS-2B cells, the intracellular BCG was significantly decreased (Fig 11A&11B and S2J&S2K Fig). However, hBD1 or/and LL37 silencing during the autophagy process weakened the autophagy function of killing the intracellular BCG (Fig 11B, 11C&11D and S2J, S2K&S2L Fig). These results indicated that hBD1 and LL37 play an important role in autophagic killing of BCG in lung epithelial cells, identifying autophagy-related AMP production as a novel mechanism of autophagy-mediated MTB killing.

**Figure 11.**
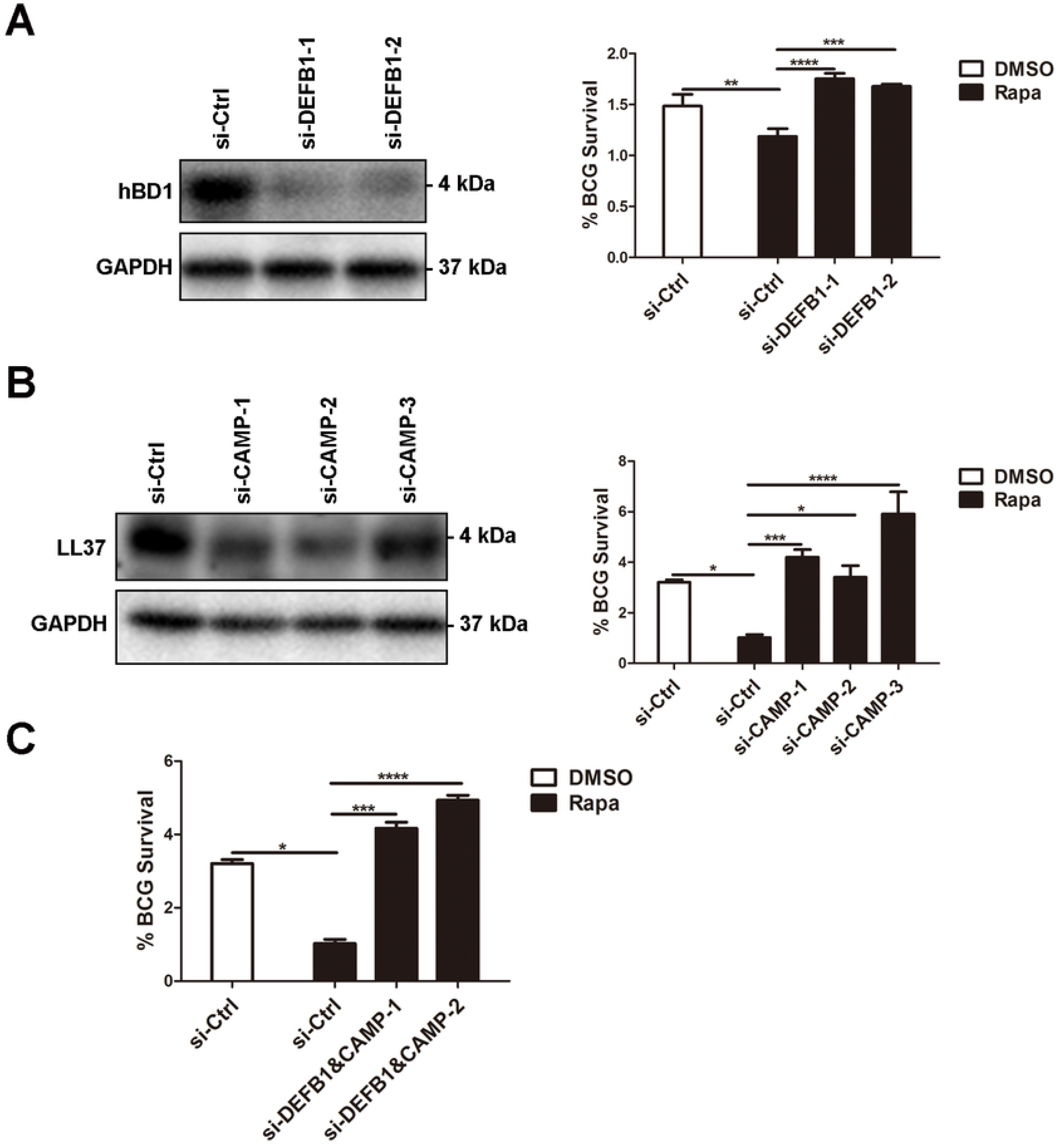
Autophagy-related production of antimicrobial peptides is a novel mechanism of autophagy-mediated BCG killing. (A-C) Silencing of hBD1 (A) or/and LL37 (B&C) weakened the autophagic killing of intracellular BCG. A549 cells were transfected with si-NC, Si-DEFB1, or/and si-CAMP for 24 h then infected with BCG (MOI = 5) for 1 h. In the rapamycin (Rapa) group the cells were pretreated by 4 μM rapamycin for 6 h prior to BCG infection. The intracellular viable bacilli were determined by CFU assays at 72 h. Survival rate was calculated compared with that at 0 h. Data are expressed as the means ± standard deviation (s.d.) \**p* <0.05, \*\**p* <0.01, \*\*\**p* <0.001, \*\*\*\**p* <0.0001. These experiments were performed independently at least thrice with similar results.

## Discussion

During pathogen invasion, AMPs can directly inhibit microbial growth and regulate inflammatory responses by attenuating damage and stimulating beneficial responses. [18]. In this study, we screened AMPs hBD1 and LL37 as highly expressing in PBMCs and alveolar epithelial cells in pathological sections of patients with TB. Furthermore, we verified that the expression of active hBD1 and LL37 involves autophagy, and that this process is related to AMP precursor cleavage with p62 also being involved. Moreover, we demonstrated that AMPs produced by autophagy could play an important role in the autophagic killing of intracellular BCG. These findings open new avenues for further research regarding anti-TB immune mechanisms and the potential for new TB treatment strategies.

To further understand the functional mechanisms of the hBD1 and LL37 in the defense against MTB invasion by lung epithelial cells, we explored the expression mechanism of hBD1 and LL37 in A549 and BEAS-2B cells. To confirm whether the AMP production process, wherein an expressed precursor form is cleaved by a proteolytic process to release the active peptide [18], is related to the occurrence of autophagy, we conducted various interventions toward the autophagy level of A549 and BEAS-2B cells. The results showed that the active AMP expression level positively correlates with the autophagy level, whereas modulating the autophagy level does not affect AMP mRNA levels. Moreover, immunofluorescence assays confirmed that the LL37 precursor hCAP18 entered the autophagosomes, revealing that the co-localization rates between hCAP18 and autophagosomes were influenced by the autophagy process in consonance with the active protein level of AMPs. These findings highlighted that the autophagic level of lung epithelial cells could affect the protein levels of hBD1 and LL37 active peptides and that this occurs in a post-transcriptional manner, likely through the autophagic cleavage of antibacterial peptide precursors. Although various previously identified AMP shearing mechanisms have been found to occur during e.g., cell exocytosis [20] and phagocytosis [32], our findings suggested that autophagy might constitute a previously uncharacterized cleavage mechanism of AMPs. In particular, whereas it was previously shown that the autophagic process can also cleave antibacterial peptide precursors [30], this cleavage mechanism had not been validated in the classical three antibacterial peptide families. Conversely, the mechanism described in the present study was also confirmed as affecting hBD1 and LL37.

We next explored the role of lysosomes in the autophagic production of hBD1 and LL37 as once the autophagosome encloses the targeted material and forms a closed bilayer membrane structure, it fuses with a lysosome and undergoes acidification to dissolve and degrade enveloped contents by forming autophagosomes [33,34]. We used Baf A1 to inhibit autophagosome and lysosome fusion to interrupt the later process of autophagy, and NH_4_Cl to prevent the acidification as a lysosomal function inhibitor, and observed whether this process affected hBD1 and LL37 production. Our analysis showed that the active protein levels of hBD1 and LL37 were down-regulated, regardless of autophagy induction by BCG or rapamycin. However, neither Baf A1 nor NH_4_Cl stimulation affected the mRNA levels of hBD1 and LL37. Additionally, these inhibitors could not affect AMP precursor hCAP18 recruitment to and entry into autolysosomes, as shown by immunofluorescence results. These results suggested that lysosome function and autolysosome formation are critical to the autophagy-related production of active hBD1 and LL37, but do not influence AMP precursor recruitment.

To evaluate how the AMP precursors selectively enter the autophagosome, we focused on the autophagic adaptor protein p62, which facilitates the function of autophagy to clear protein aggregates. In neurodegenerative diseases, p62 can mediate the transport of cytosolic proteins to lysosomes and effect their cytoplasmic proteolysis into AMPs in macrophages [35]. Notably, this function appears to be specific to p62, as other autophagic adaptor proteins, such as NBR1, do not exhibit the ability to recruit MTB phagosomes in the autophagy process, although NBR1 in particular shows similar function to p62 during its interaction with LC3 and recruitment of ubiquitinated proteins [35–37]. Additional studies revealed that lysosomal hydrolase function and the acidification of MTB-containing vesicles are completely independent of p62 in autophagy up-regulated macrophages, whereas p62 plays a special role in autophagic antimicrobial activity [30].

Therefore, we interrogated whether p62 is involved in the production of hBD1 and LL37 active peptides. Our results showed that silencing p62 in A549 cells down-regulated hBD1 and LL37 active protein levels but did not affect their mRNA levels. Moreover, co-localization was observed among p62, hCAP18, autophagosomes, and lysosomes with p62 able to directly interact with hCAP18, as shown by immunofluorescence and immunoprecipitation, respectively. These results suggested that p62 is involved in active hBD1 and LL37 production, with the direct interaction with hCAP18 indicating that p62 affects AMP expression by recruiting antimicrobial peptide precursors to autophagosomes.

Autophagy can play a role in fighting MTB infection in the natural immune process through a variety of mechanisms, such as facilitating the resolution of intracellular MTB, producing functional cytokines and effector proteins, or regulating inflammatory responses [38–40]. Based on the findings of the present study, we hypothesized that autophagy produces active AMPs to exert its antibacterial function against intracellular MTB. To verify this conjecture, we performed a colony formation assay. Statistical analysis of the survival rate of BCG in A549 and BEAS-2B cells showed that following rapamycin treatment, the intracellular BCG survival rate decreased, whereas this effect was ameliorated upon concurrent silencing of hBD1 and/or LL37. This phenomenon indicated that the AMP production plays a crucial role in the process of anti-intracellular BCG infection in autophagy-up-regulated lung epithelial cells, highlighting AMP production as an important mechanism by which autophagy plays its role in killing MTB.

In summary, we observed the high expression of hBD1 and LL37 in PBMCs and lung epithelial type II cells of patients with TB and in A549 and BEAS-2B cells infected with BCG, and verified the involvement of autophagy and autolysosomes in the production of active hBD1 and LL37, and of autophagic adaptor protein p62 in the selective recruitment of AMP precursors. Our results suggested that classic AMP precursors could be sheared during the autophagy process, thus producing active AMPs to exhibit anti-MTB activity. These findings will enhance our understanding regarding the antibacterial function of epithelial cells in the process of natural immunity during MTB invasion, and provide a new perspective for designing new treatments for TB.

## Availability of data and materials

The datasets used and/or analyzed during the current study are available from the corresponding author on reasonable request.

## Ethics approval and consent to participate

Prior to sample collection, written informed consents were obtained and the study approved by the Ethics Committee of the Southern Medical University.

## Consent for publication

All healthy volunteers provided informed consent and agreed to the publication of relevant data and related images.

## Supporting information

**Supplementary Figure S1. Electron microscopy analysis of BCG-infected lung epithelial cells.**

A549 and BEAS-2B cells were infected with BCG for 1 h (MOI = 5). The bacilli were observed inside cells (red arrow) and enveloped by the bilayer membrane autophagosomes (yellow arrow).

**Supplementary Figure S2. Antimicrobial peptides hBD1 and LL37 were highly expressed in BCG-infected lung epithelial cells and exhibit anti-MTB activity.**

The active peptides of hBD1 and LL37 were detected at indicated time points and evaluated by western blotting. (B, C) *hBD1* (b) and *LL37* (C) mRNA expression in BCG-infected BEAS-2B cells was detected at indicated time points using real-time PCR. (D) The autophagic level of BEAS-2B cells affected the active peptide levels of hBD1 and LL37. BEAS-2B cells were pretreated with 4 μM rapamycin for 6 h and 5 mM 3-MA for 2 h and then infected with BCG for 24 h. The active peptide levels of hBD1 and LL37 were evaluated using western blotting. (E, F) The autophagic level of BEAS-2B cells did not affect the mRNA level of hBD1 (E) and LL37 (F). BEAS-2B cells were pretreated and infected with BCG as described above. The mRNA levels of hBD1 and LL37 were evaluated using real-time PCR. (G-I) Silencing of hBD1 or/and LL37 decreased intracellular BCG killing in BEAS-2B cells. The intracellular viable bacilli were determined using a CFU assay at 72 h. The survival rate of BCG was calculated in comparison with that at 0 h. (J-L) Silencing of hBD1 or/and LL37 weakened the autophagic killing of intracellular BCG. BEAS-2B cells were transfected with si-NC, si-DEFB1, or/and si-CAMP for 24 h, then infected with BCG (MOI = 5) for 1 h. In the rapamycin (Rapa) group, the cells were pretreated with 4 μM rapamycin for 6 h prior to BCG infection. The intracellular viable bacilli were determined using CFU assays at 72 h. Survival rate was calculated compared with that at 0 h. Data are expressed as the means ± standard deviation (s.d.). \**p* < 0.05, \*\**p* < 0.01, \*\*\**p* < 0.001, \*\*\*\**p* < 0.0001. These experiments were performed independently at least thrice with similar results.

**Supplementary Figure S3. LL37 expression level in various TB lesions.**

High level expression of LL37 was detected in human alveolar type II pneumocytes from patients with TB via immunohistochemistry. Micrograph shows strong LL37 immunostaining in human alveolar type II pneumocytes from patients with TB compared to that in granuloma, caseous necrosis, and immune cells (magnification ×200). (B) Average optical of immunohistochemistry photographs from (A). Data are expressed as the means ± standard deviation (s.d.). \*\*\*\**p* < 0.0001. These experiments were performed independently at least thrice with similar results.

**Supplementary Figure S4. AMP expression levels in various cell types.**

A549, BEAS-2B, THP1, and HMDMs were infected with BCG for 48 h and the hBD1 and LL37 mRNA were detected using qPCR. Data are expressed as the means ± standard deviation (s.d.). \**p* < 0.05, \*\**p* < 0.01, \*\*\*\**p* < 0.001. These experiments were performed independently at least thrice with similar results.

## References

1. World Health Organization. Global Tuberculosis Report 2019. TB Fact sheet [Internet]. 2019;1–2. Available from: https://apps.who.int/iris/bitstream/handle/10665/329368/9789241565714-eng.pdf?ua=1

2. Bermudez LE, Goodman J. Mycobacterium tuberculosis invades and replicates within type II alveolar cells. Infect. Immun. 1996;64:1400–1406.

3. Fine KL, Metcalfe MG, White E, et al. Involvement of the autophagy pathway in trafficking of Mycobacterium tuberculosis bacilli through cultured human type II epithelial cells. Cell. Microbiol. 2012;14:1402–1414.

4. Mehta PK, King CH, White EH, et al. Comparison of in vitro models for the study of Mycobacterium tuberculosis invasion and intracellular replication. Infect. Immun. 1996;64:2673–2679.

5. Dobos KM, Spotts EA, Quinn FD, et al. Necrosis of lung epithelial cells during infection with Mycobacterium tuberculosis is preceded by cell permeation. Infect. Immun. 2000;68:6300–6310.

6. Bevins CL, Salzman NH. Paneth cells, antimicrobial peptides and maintenance of intestinal homeostasis. Nat. Rev. Microbiol. 2011. p. 356–368.

7. Harder J, Meyer-Hoffert U, Teran LM, et al. Mucoid Pseudomonas aeruginosa, TNF-alpha, and IL-1beta, but not IL-6, induce human beta-defensin-2 in respiratory epithelia. Am. J. Respir. Cell Mol. Biol. 2000;22:714–721.

8. Mendez-Samperio P, Miranda E, Trejo A. Mycobacterium bovis Bacillus Calmette-Guerin (BCG) stimulates human beta-defensin-2 gene transcription in human epithelial cells. Cell Immunol [Internet]. 2006;239:61–66. Available from: http://www.ncbi.nlm.nih.gov/entrez/query.fcgi?cmd=Retrieve&db=PubMed&dopt=Citation&list_uids=16762333.

9. Yang X, Cheng YT, Tan MF, et al. Overexpression of porcine beta-defensin 2 enhances resistance to Actinobacillus pleuropneumoniae infection in pigs. Infect. Immun. 2015;83:2836–2843.

10. Jarczak J, Kościuczuk EM, Lisowski P, et al. Defensins: Natural component of human innate immunity. Hum. Immunol. 2013. p. 1069–1079.

11. Castaneda-Sanchez JI, Garcia-Perez BE, Munoz-Duarte a R, et al. Defensin production by human limbo-corneal fibroblasts infected with mycobacteria. Pathogens [Internet]. 2013;2:13–32. Available from: http://www.embase.com/search/results?subaction=viewrecord&from=export&id=L372464405 http://dx.doi.org/10.3390/pathogens2010013.

12. García-Pérez BE, Villagómez-Palatto DA, Castañeda-Sánchez JI, et al. Innate response of human endothelial cells infected with mycobacteria. Immunobiology. 2011;216:925–935.

13. Ganz T. Antimicrobial polypeptides in host defense of the respiratory tract. J. Clin. Invest. 2002. p. 693–697.

14. Schutte BC, McCray PB. [Beta]-Defensins in Lung Host Defense. Annu. Rev. Physiol. [Internet]. 2002;64:709–748. Available from: http://www.ncbi.nlm.nih.gov/pubmed/11826286.

15. Ganz T. Defensins: Antimicrobial peptides of innate immunity. Nat. Rev. Immunol. 2003. p. 710–720.

16. Tecle T, Tripathi S, Hartshorn KL. Defensins and cathelicidins in lung immunity. Innate Immun. 2010. p. 151–159

17. Martineau AR, Newton SM, Wilkinson KA, et al. Neutrophil-mediated innate immune resistance to mycobacteria. J. Clin. Invest. 2007;117:1988–1994.

18. Lai Y, Gallo RL. AMPed up immunity: how antimicrobial peptides have multiple roles in immune defense. Trends Immunol. 2009. p. 131–141.

19. Castaneda-Sanchez JI, Garcia-Perez BE, Munoz-Duarte a R, et al. Defensin production by human limbo-corneal fibroblasts infected with mycobacteria. Pathogens [Internet]. 2013;2:13–32. Available from: http://www.embase.com/search/results?subaction=viewrecord&from=export&id=L372464405 http://dx.doi.org/10.3390/pathogens2010013.

20. Sørensen OE, Follin P, Johnsen AH, et al. Human cathelicidin, hCAP-18, is processed to the antimicrobial peptide LL-37 by extracellular cleavage with proteinase 3. Blood. 2001;97:3951–3959.

21. Valore EV, Ganz T. Posttranslational processing of defensins in immature human myeloid cells. Blood [Internet]. 1992;79:1538–1544. Available from: http://www.ncbi.nlm.nih.gov/pubmed/1339298.

22. Yount NY, Wang MS, Yuan J, et al. Rat neutrophil defensins. Precursor structures and expression during neutrophilic myelopoiesis. J. Immunol. 1995;155:4476–4484.

23. Rice W, Ganz T, Kinkade J, et al. Defensin-rich dense granules of human neutrophils. Blood. 1987;70:757–765.

24. Di Nardo A, Vitiello A, Gallo RL. Cutting Edge: Mast Cell Antimicrobial Activity Is Mediated by Expression of Cathelicidin Antimicrobial Peptide. J. Immunol. [Internet]. 2003;170:2274–2278. Available from: http://www.jimmunol.org/cgi/doi/10.4049/jimmunol.170.5.2274.

25. Nell MJ, Sandra Tjabringa G, Vonk MJ, et al. Bacterial products increase expression of the human cathelicidin hCAP-18/LL-37 in cultured human sinus epithelial cells. FEMS Immunol. Med. Microbiol. 2004;42:225–231.

26. Gutierrez MG, Master SS, Singh SB, et al. Autophagy is a defense mechanism inhibiting BCG and Mycobacterium tuberculosis survival in infected macrophages. Cell. 2004;119:753–766.

27. Ponpuak M, Davis AS, Roberts EA, et al. Delivery of Cytosolic Components by Autophagic Adaptor Protein p62 Endows Autophagosomes with Unique Antimicrobial Properties. Immunity. 2010;32:329–341.

28. Singh SB, Davis AS, Taylor GA, et al. Human IRGM induces autophagy to eliminate intracellular mycobacteria. Science (80-.). 2006;313:1438–1441.

29. Jo EK. Autophagy as an innate defense against mycobacteria. Pathog. Dis. 2013. p. 108–118.

30. Davis AS, Roberts EA, et al. Delivery of Cytosolic Components by Autophagic Adaptor Protein p62 Endows Autophagosomes with Unique Antimicrobial Properties. Immunity. 2010;32:329–341.

31. Alonso S, Pethe K, Russell DG, et al. Lysosomal killing of Mycobacterium mediated by ubiquitin-derived peptides is enhanced by autophagy. Proc. Natl. Acad. Sci. [Internet]. 2007;104:6031–6036. Available from: http://www.pnas.org/cgi/doi/10.1073/pnas.0700036104.

32. c, Yamasaki K, Kabigting FD, et al. Kallikrein expression and cathelicidin processing are independently controlled in keratinocytes by calcium, vitamin D 3, and retinoic acid. J. Invest. Dermatol. 2010;130:1297–1306.

33. Mizushima N, Levine B, Cuervo AM, et al. Autophagy fights disease through cellular self-digestion. Nature. 2008. p. 1069–1075.

34. Xie Z, Klionsky DJ. Autophagosome formation: Core machinery and adaptations. Nat. Cell Biol. 2007. p. 1102–1109.

35. Bjørkøy G, Lamark T, Brech A, et al. p62/SQSTM1 forms protein aggregates degraded by autophagy and has a protective effect on huntingtin-induced cell death. J. Cell Biol. 2005;171:603–614.

36. Pankiv S, Clausen TH, Lamark T, et al. p62/SQSTM1 binds directly to Atg8/LC3 to facilitate degradation of ubiquitinated protein aggregates by autophagy*[S]. J. Biol. Chem. 2007;282:24131–24145.

37. Kirkin V, Lamark T, Sou YS, et al. A Role for NBR1 in Autophagosomal Degradation of Ubiquitinated Substrates. Mol. Cell. 2009;33:505–516.

38. Klionsky DJ, Emr SD. Autophagy as a regulated pathway of cellular degradation. Science (80-.). 2000. p. 1717–1721.

39. Sultana Rekha R, Rao Muvva SJ, Wan M, et al. Phenylbutyrate induces LL-37-dependent autophagy and intracellular killing of mycobacterium tuberculosis in human macrophages. Autophagy. 2015;11:1688–1699.

40. Shin DM, Yuk JM, Lee HM, et al. Mycobacterial lipoprotein activates autophagy via TLR2/1/CD14 and a functional vitamin D receptor signalling. Cell. Microbiol. 2010;12:1648–1665.

